# Deep Recurrent Neural Network and Point Process Filter Approaches in Multidimensional Neural Decoding Problems

**DOI:** 10.1101/2020.08.10.244368

**Authors:** Mohammad Reza Rezaei, Behzad Nazari, Saeid Sadri, Ali Yousefi

## Abstract

Recent technological and experimental advances in recording from neural systems have led to a significant increase in the type and volume of data being collected in neuroscience experiments. This brings an increasing demand for development of appropriate analytical tools to analyze large scale neuroscience data. Simultaneously, advancement in deep neural networks (DNNs) and statistical modeling frameworks have provided new techniques for analysis of diverse forms of neuroscience data. DNNs like Long short-term memory (LSTM) or statistical modeling approaches like state-space point-process (SSPP) are widely used in the analysis of neural data including neural coding and inference analysis. Despite wide utilization of these techniques, there is a lack of comprehensive studies which systematically assess attributes of LSTM and SSPP approaches on a common neuroscience data analysis problem. As a result, this occasionally leads to inconsistent and divergent conclusions on the strength or weakness of either of the methodologies and also statistical significance of the analytical outcomes. In this research, we focus on providing a more systematic and multifaceted assessment of LSTM and SSPP techniques in a neural decoding problem. We examine different settings and modeling specifications to attain the optimal modeling solutions. We propose new LSTM network topologies and approximate filter solution to estimate a rat movement trajectory in a 2-D spaces using an ensemble of place cells’ spiking activity. For each technique; we then study performance, computational efficiency, and generalizability of each technique in this decoding problem. By utilizing these results, we provided a succinct picture of the strength and weakness of each modeling approach and suggest who each of these techniques can be properly utilized in neural decoding problems.

## 1 Introduction

Recent advancements in sensor technology and progresses in clinical experiments have provided ability to simultaneously record neural activity of different brain areas in human and non-human primates [1–3]. Analyzing these data will help us to understand what stimuli or event elicits a particular pattern of neural activity [4–6] or how complex processes in brain, like learning [7] and memory, are shaped [8]. Higher dimension and volume of the neural data recorded in these experiments present a unique set of challenges and difficulties in their analysis and have led to an elevated demand for more accurate and computationally efficient solutions [9–11]. In this research, we focus on a neural decoding problem which is based on processing spiking data recorded from a cell ensemble representing an external behavioral or physiological correlates like movement. We utilize deep neural network (DNN) and state-space (SS) modeling frameworks to process the neural data and decode relevant behavioral correlates.

SS have provided powerful tools in analysis of neuroscience data [12–14]. For instance, state-space poin-process (SSPP) has been successfully applied to estimate a rat position in 2-D spaces from hippocampus place cells’ spiking activity [15], tracking oscillatory activities in the subthalamic nucleus of Parkinson’s patients [16], and decoding arm movement and grasp actions from neural activity of motor cortex area [13]. In parallel, deep neural networks (DNNs) have been utilized in neuroscience data analysis to address a similar class of decoding and inference problems. For example, Long Short-term Memory (LSTM) [17], a particular recurrent neural networks (RNN) [18], is used to charactrize direct input and output relationship between neural activity and physilogical variables such as decoding from motor and somatosensory cortices [19], Self-localization in 2-D spaces [20]. It also used as decoder model in encoding-decoding problems like decoding from human motor cortical signals [21], and decoding a rat position in 2-D spaces using hippocampus place cells spiking activity [22].

Despite the extensive utilization of DNN and SS approaches in analysis of neuroscience data, there are still multiple modeling and computational challenges to be addressed. In SSPP approach, the neural spiking activity are formulated as a function of underlying latent dynamical states, and dynamical state models are defined by a state-space point process. The established solution for SSPP uses recursive Bayes filter which consists of prediction and update rules defined by Chapman-Kolmogorov and Likelihood function. The solution is appropriate for a one-dimensional decoding problem, but becomes computationally expensive as the dimension of the decoding problem grows; even a 2-D decoding problem becomes computationally demanding [23]. Tools and techniques like Gaussian approximation [24, 25], sequential Monte-Carlo (SMC) methods including particle filtering [26, 27] have been applied to address computational challenge. However, these solutions might be less efficient in multi-dimensional decoding problems.

DNNs also are not immune to the computational complexcity. DNNs are data greedy; the number of the model’s parameters are thousands of times larger than the traditional neural networks like shallow neural network and they demand large datasets for a proper traning [28]. When a DNN is trained with a small dataset, it simply overfits to the training data and lose its accuracy and generalizability on the test data [29]. Even though this problems are partially addressed using synthetic training data [26] or dropout technique [29], there are ongoing efforts to reduce the size (number of parameters) of DNNs. In practice, this is a challenging task as construction of an efficient network for a specific problem requires a lot of experience and good insight of the problem in hand [30]. The other issue with DNNs is the lack of model interpretabilty, which is a major caveat of DNNs [31]. A remedy to overcome this problem which is inspired from biological neural system is hierarchical processing of features by increasing level of model interpretability in deeper layers of neural systems [32]. By embedding this charactrestic in DNN, the network accuracy and generalizability showed a significanct boost [33, 34]. Even though DNNs are used for neural decoding problems, but none of them addressed interpretation ability of DNNs for neural decoding problems [19, 20]. So, building interpretable DNNs for neuroscience data which have complex dynamics is still a modeling challenge and need to be addressed to elevate potential of DNNs for neural decoding problems.

Here, we focus on decoding a rat movement trajectory in 2-D spaces from neural activity recorded from an ensemble of place cells. Through this problem, we aim to revisit and potentially address these challenges. Despite established solutions [12] for 2-D decoding problems, this problem has room to be better addressed, particularly when the experiment is being conducted in a complex maze structure [23]. We address this problem by proposing DNN and SSPP models to estimate the rat movement. We work to assess different aspects of both technqiues including accuracy, computational efficiency, and generalizability, which are crucial to build a succint picture of each methodology’s strength and weaknesses.

For SSPP, we examine Numerical exact solution and an approximate filter solution. Numerical exact solution, uses a Bayes filter solution [28]. The posterior distribution of the rat position in the maze is estimated by solving the filter update and likelihood functions numerically. For the approximate solution, we propose a approximate filter solution with a drop-merger algorithm proposed in [35] to estimate the rat movement trajectory. For DNN approach, we propose LSTM network topologies to derive an architecture that properly captures structural dynamics of the rat’s movement by presenting interpretable elements of movement from its spiking activity in a hierarichal structure. This research provides a new set of modeling approaches in decoding from neural activity problems and helps to reach a more clear picture of pros and cons of DNN and SSPP in neural decoding problems and eventually provides alternative choices in this field.

In the next sections, we discuss our proposed SSPP models and LSTM network topologies to estimate a rat movement trajectory in 2-D spaces (W-shaped maze) in details. We then discuss the 2-D decoding problem and the dataset being used in this research. We run different tests to check performance, computational complexity and generalizability of both approaches. Finally, we investigate how the relationship between place cells and rat position is being captured in them. Using this result, we eventually provides alternative choices in neural decoding problems.

## 2 Methods

In this section, we describe SSPP and LSTM modeling approaches to decode 2-D movement trajectory from an ensemble spiking activity and explain the theory behind each technique. For SSPP, we define how the relationship between the movement trajectory and spiking activity can be formulated. We build the state and observation processes’ equations and define the relationship between the state variables and observed neural activity. For LSTMs, we represent an hirariacal structure in LSTM network topology. We incorporate higher information of the rat movement including geometry of the maze, the movement velocity, and the movement direction into the network topology to increase accuracy and interpretability of the network topology. We discuss how the decoding accuracy improves by incorporating these information in a hiarical structure in the network topology.

### 2.1 State-space point-process approach

Here, we discuss two solutions: Numerical exact solution and an approximate filter solution. The first solution, Numerical exact solution, uses a Bayes filter solution [36]. The posterior distribution of the rat position in the maze is estimated by solving the filter update and likelihood functions numerically. In this solution, each cell firing activity is modeled by a conditional intensity function (CIF) which is built using a non-parametric method to characterize the spiking activity of individual neurons as a function of the rat position in a 2-D maze [23]. For the second solution, which is an approximate filter solution, we assume that the posterior distribution of the rat position per each time-step can be approximated using a Gaussian Mixture Model (GMM). In this solution, each cell’s CIF is defined by Mixture of Gaussian (MoG). Utilizing this specific form of CIF function, we build a computationally efficient algorithm that estimates the optimal number of mixture components and corresponding parameters per each time-step given the observed spiking activity [35]. This solution is computationally efficent in 2-D and higher-dimensional decoding problems where the computation of numerical solution becomes demanding [37]. In the next part we describe each model in detail.

### 2.1.1 Exact solution

For a point-process observation, we model spiking activity of a neuron by representing its firing rate as a function of neural correlates such as position plus the history of the population spiking activity. The firing rate of a neuron at time *t* is defined by

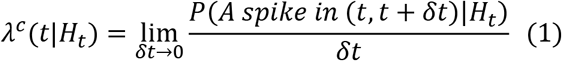

where, *c* is the neuron index and *H_t_* represents the full history of spiking for all neurons up to time *t* [38, 39]. For the specific problem of this research, we use a non-parametric kernel method to build each neuron’s firing rate - or CIF – as a function of *X_k_* = (*x_k_*, *y_k_*), where (*x_k_*, *y_k_*) represent the rat’s position at the time index *k* in a 2-D spaces [40]. Given the neural ensemble spiking activity, the likelihood that the rat is at coordinate *X_k_* given observing Δ*N_k_* total spikes from an ensemble of *C* cells, in the interval *Δ_k_* = (*t_k_−1*, *t_k_*] is defined by

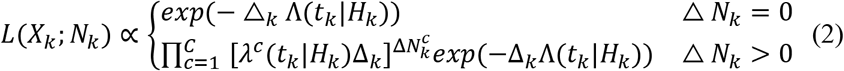

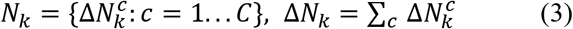

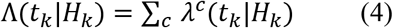

where *c* refers to the cell index and λ^*c*^(*t_k_*|*H_k_*) represents the *c*^*th*^ cell intensity as a function of *X_k_*, and Δ*k* defines the time-step. We assume that the rat movement over the 2-D spaces follows a Markov process, which is defined by

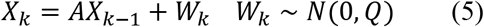

where, *A* is the state matrix, *W* is the process noise, and *Q* is the process noise covariance matrix. Under this assumption, the one-step density of state *X_k_*, is defined by

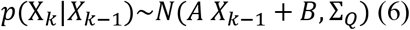

Using equations (2) and (5), we can run the recursive Bayes filter [13] to estimate the rat movement using the spiking activity of the place cell ensemble. The likelihood function does not follow a Gaussian distribution, and thus the filter solution does not have an analytical solution. Base on [41], the posterior distribution of the state at time index *k* can be derived by

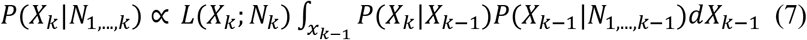

where the integral over *X_k−1_* defines the one-step prediction in the Bayes filter paradigm [36]. The term *P*(*X_k_*|*N*_1_…*k*) is the filter estimation at time index *k* given {*N*_1_, …, *N_k_*}. The computational complexity of the exact solution grows exponentially as the dimension of the decoding problem increases [23]. As a result, for a 2-D decoding problem the computational cost becomes prohibitive; especially when the filter is required to be estimated with a fine spatial resultion in each axis of X – here, x and y. In the following section, we discuss a computationally efficient approximate solution which can be used for real-time decoding problems in multi-dimensional spaces.

### 2.1.2 Approximate filter solution

There are approximate solutions like Gaussian approximataion solutions with a low computational cost that use a Gaussian approximation of the posterior distribution; however, multi-modality of cells’ receptive field make this solution less accurate in decoding the rat position as the number of cells in the recording grows. We propose a new solution, where filter is approximated by GMM and each neuron’s CIF is defined by a MoG. The CIF for a neuron is defined by

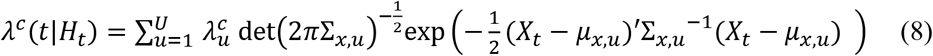

where, 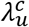 defines the rate of *u^th^* mixture model related to *c^th^* cell. *μ_x,u_* and *Σ_x,u_* are the mean and covariance of the *u^th^* mixture model over *X_t_* space. For each cell, there are U mixture components, where the number of mixture component for a neuron may differ from other neurons. It is worth to mention that sum of *λ_u_* is not normalized and it can be any positive number depending the cell firing rate over the space. The likelihood of the rat position given the ensemble spiking activity is the same as the one defined in equation (2), and the rat movement trajectory over the space is defined by equation (5). In this filter solution, we assume that the posterior distrubtuin of the rat position in the space has multi-modal distribution which can be approximated by a GMM,

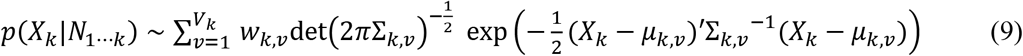

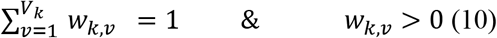

where *V_k_* is the number of mixture components at time index *k* and *w_k,v_* is the weight of the *v^th^* mixture component with (*μ_k,v_*, *Σ_k,v_*) paramaters. Appendix A describes how each of these parameters is estimated over different time points. Computation time of this filter solution does not grow exponentially with the decoding problem dimension, and as a result, it can be applied to high dimensional decoding problems without an exponential growth in the computational cost [23]. Note that the number of mixture components on each processing time can be optimally controlled; thus, its computational time can be adjusted depending on the processing time and accuracy requirements [35].

In this section, we introduced both exact and the approximate filter solutions for the 2-D decoding problem. We revisit these techniques and study their performance and computational efficiency along with DNNs.

### 2.2 Deep neural network approach (LSTM network topologies)

In this section, we argue that a directly mapping the input – spiking activity to the rat position is not necessary the best toplogy from both interprtatbility and prediction accuracy viewpoints. Then, we propose an hierarchal architecture that properly captures structural dynamics of the rat’s movement by extracting higher order elements of its movement in a hierarchal structure. We suggest this hierarchal LSTM network toplogy along with a proper training constraints not only reaches a higher prediction accuracy, but also captures higher elements of movements in the enural activity.

### 2.2.1 Higher-order movement components and how to decode them in LSTM network

a rat movement trajectory depends on the geometry of the area that rat moves in, the rat velocity – and in particular, the direction of its movement. These provides higher-order information about the rat movement and its position. As a result, embedding these information as a part of LSTM toplogy might improve accuracy of the rat movement estimation. However, the key is how different layers of the LSTM are constuctued and communication with each other. We propose a hierchical structure where deeper layers capture higher-order information of the rat movement and uses them to estimate the rat position in the maze. We propose two decoding steps (layers): decoding the maze toplogy by segementing the experiment area to subregions and decoding derivations of movement. We use them to improve accuracy of the rat movement estimation.

The movement constraints by the shape of the maze and walls inside and outside area of the maze. So, we can consider the geometrical features as areas that a rat can – alos cannot - move in it. There are multiple approaches that we can embed this information in a DNN [42]. The approach we utilize, segments the experiment area to multiple regions (segments). By applying this segmentation method and a Multiple Logistic Regression (MLR) [43], we can estimate the probability of rat being in each of these segmenet given the ensemble spikign activity. Assume that the maze structure is a W-shaped, **Figure 1.a**. We want to segment its area for encoding model. The first strategy that reaches to mind is to segment the whole area of the experiment to equal segments, like **Figure 1.b**. In this strategy the area outside the maze is segmented to multiple segments, but it is not necessary. Because the rat won’t walk through the outside area at all and information of exact locations outside the maze is not useful, so we can model it as a one segment and reduce number of segments, **Figure 1.c**. The probability information of this area can be interperate as a penalty term for the position decoder to estimate positions outside the maze.

**Figure 1:**
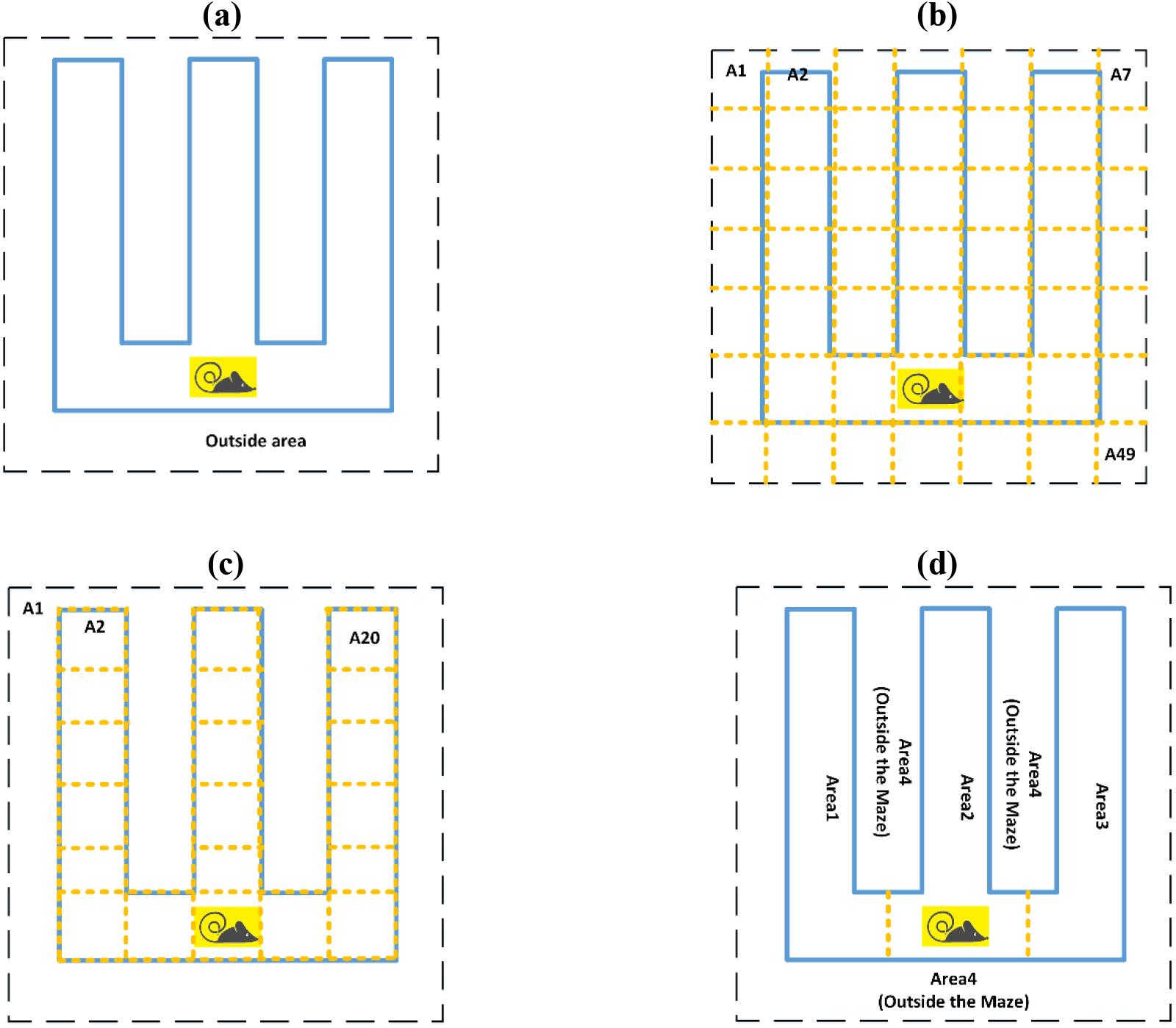
Different mechanisms to encode the geometry of the maze. **a**. The experiment maze structure, where the rat moves in, blue lines represents the walls and borders of the maze; and the black lines are boards of the experiment area. **b.** The maze area is segmented using multiple small squares. Here, we have 49 different segments representing the whole area including both inside and outside maze. **c.** The experiment area divided one segment covers outside area and 20 segments covers inside the maze **d**. A balanced segmentation of the maze topology, the maze area is encoded by three segments, and one segment for the outside area.

In addition, the area inside of the maze, by considering the maze shape (the maze has three arms), can be divided to three general segments (number of arms) like **Figure 1.d**. This can reduce the number of segment and still convey information of moving path between arms of the maze. This is the optimum way to segment inside area of the maze, because If we reduce this number we cant extract enough information. For example if we consider one segment for the inside are, we can’t gain anything because we already have this information by using information of the outside area. The rat always can move from Area-1 to Area-2 and it can’t be in Area-1 and in the next time-step jumps to Area-3 without passing Area-2, so this can be interpreted as continuity information. This segmentation method provides penalty term for the position-estimator layer to avoid estimating wrong positions outside the experiment structure area by providing information of rat position in the area-4. And provides continuity term for movement inside the maze, by providing information of transition between the inside areas, area-1 to area-3.

For decoding information about derivations of the movement from neural activity we can use a regressor layer for each derivation order. This regressor layer could be any type of linear or non-linear regressors. For example, to extract information about the first derivation of movement (rat’s velocity) from neural activity, we can build the regressor layer by two LSTM units (because we have 2-D velocity). Each unit extract velocity information for a specific axis (*x* and *y* axis). This information can helps the position estimator layer to estimate the 2-D position with considering first order dynamics of the rats’ movement.

### 2.2.2 LSTM network toplogies deriven by high-order movement terms

To have a refrence model to assess possible improment in prediction result, we use a traditional LSTM topology (the 1^st^ LSTM topology) with formulation described in [44]. This topology has been used widely to decode neural spiking activity [19, 20, 22, 44] and it is our reference model to compare prediction accuracy and interpretability of network topologies being proposed in this research. **Figure 2** shows the block diagram of the 1^st^ LSTM model; in this network topology, higher-order terms of the movement like velocity or geometry decoded at the same order without considering any hierarchal orders among the network layers. Based on physical rules of movement, these information improves prediction accuracy if being used in a proper and coronoligcal order. So, the key to use them is in what order these information should extract in the network. In the this section, we discuss the hierarchal topologies we developed to address this challenge.

**Figure 2:**
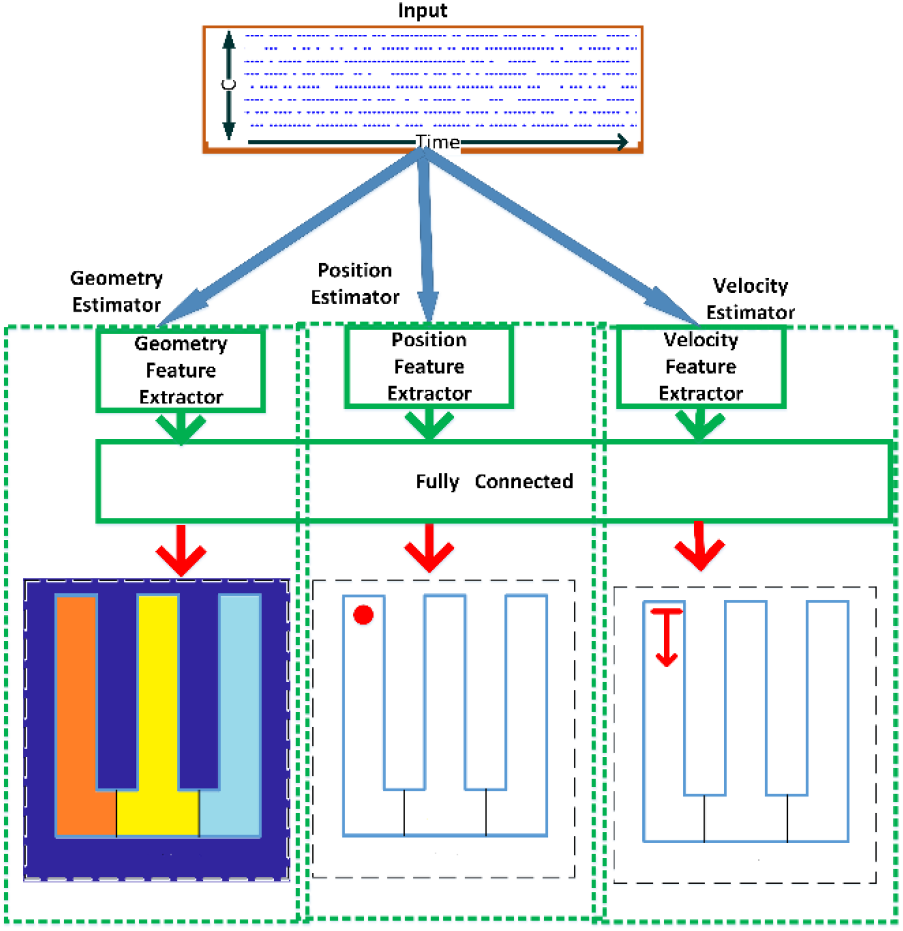
Traditional LSTM network topology (the 1^st^ model). This model consists of a layer of LSTM recurrent units and a fully connected output layer to estimate the rat position -(*x*, *y*) - from the place cells’ ensemble spiking activity. The input is spiking activity of an ensemble of C place cells and the output is −2D position of the rat in the maze (identified by a red dot indicator in the output image). Red segment in geometry-estimator section is the segment with highest probability where the rat is in it in the current time step. Finially, red arrow in velocity estimator section is direction of the rat movement.

We propose our 2^nd^ LSTM network topology with the ability to extract information about the geometrical features (mentioned in previous section) of the area that a rat moves in, prior to estimation of the rat position. The “position-estimator” layers uses this higher order movement term to estimate the rat’s position - shown in **Figure 3.a**. In this topology, the second layer of the network, called “geometry-estimator” layer, extracts the geometrical features from the cell ensemble neural activity and passes it to the output layer, position-estimator layer. This extra information which is being passed to the output layer improves the decoding accuracy in predicting the rat position if our assumption mentioned in previous section and proposed hierarchy is correct. This is because if the geometry layer does not carry information about the position, we then expect the prediction performance not to significantly gorw compared to 1^st^ LSTM. Also this geometry-estimator layer restricts the output of the position estimator from estimating positions outside the maze area (see previous section) and leads to higher position estimation accuracy.

**Figure 3:**
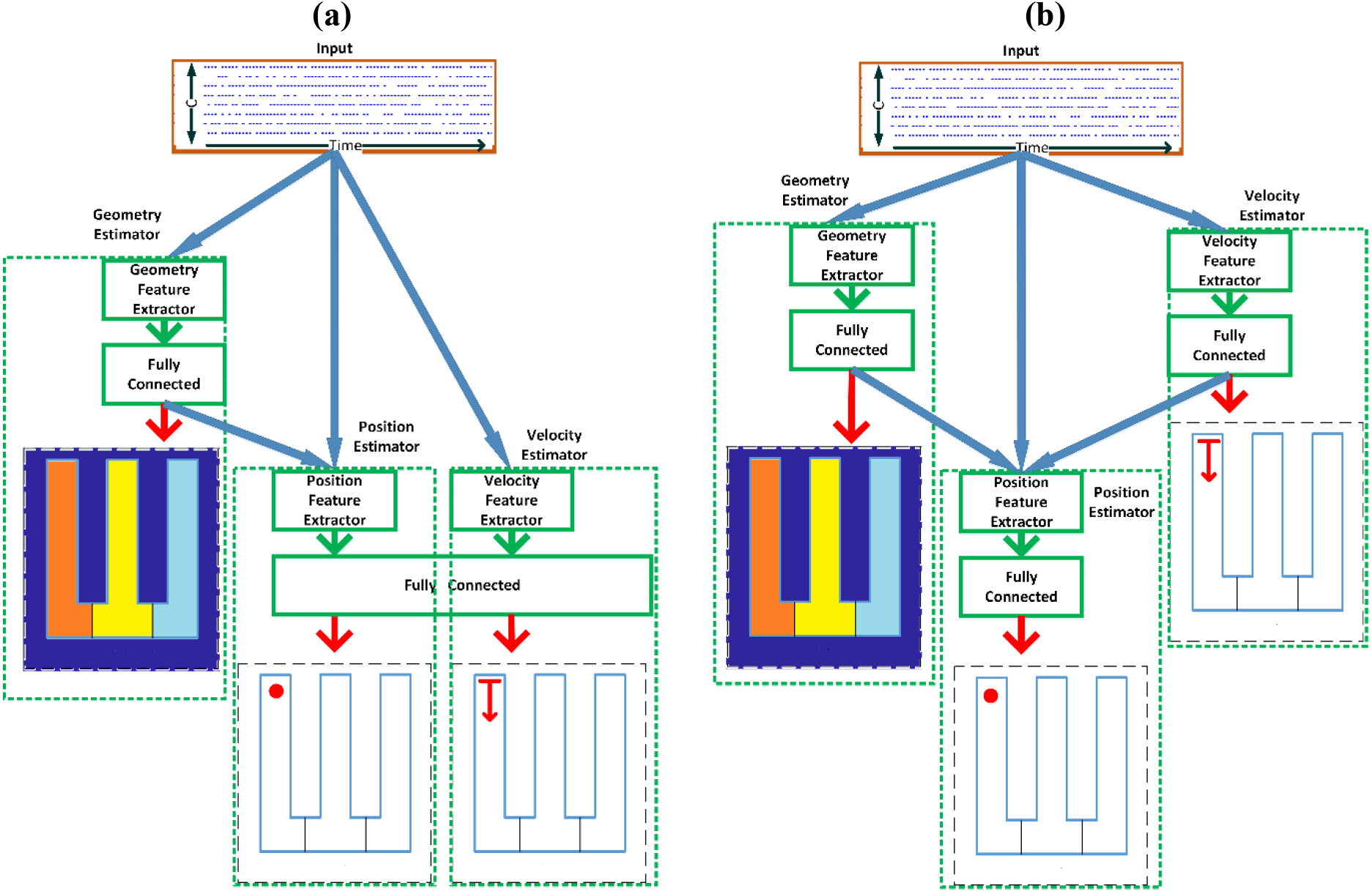
The propsed LSTM network topologies. **a**. The 2^nd^ LSTM network topology. Similar structure to the 1^st^ topology for position estimator with addition of the geometry-estimator layer that adds information about geometry of the environment as an input to the output layer. Green boxes imaginary separate the layers and visualization of the layers output. The output of geometry-estimator layer shows that the area that the rat is at the current time step, area-1 (left arm of the maze identified by red color). **d**. The 3^rd^ LSTM network topology. In this topology, the movement velocity (red arrow) is extracted directly from spiking activity and passed to the position-estimator layer along with geometry information.

The 3^rd^ LSTM network topology is simialrt to the 2^nd^ LSTM network topology, but instead of estimating position and velocity in a same order, it extracts velocity features which are the derivative of position as prior knowledge for estiamtiong the position and pass this higher order movement term to the position-estimator layer to estimate the rat’s position; **Figure 3.b** shows our 3^rd^ LSTM network topology. In this topology, the “velocity-estimator” layer first decodes information about first order dynamics of the rat’s movement using LSTM units and passes decoding result to the position-estimator layer to estimate the rat position. Because the velocity features are not related to the geometrical features we can extract them in the same order (prior to the position-estimator layer) with geometrical features, but independently. Like before this extra information which being passed to the output layer, improves the decoding accuracy in predicting the rat position if our assumption and proposed hierarchy is correct.

### 2.2.3 Cost function and training constraints

In neuroscience experiments that involve studying a rat movement in a maze, the rat spends a significant amount of time on places which it gets rewards or needs a turnaround; however, the spiking activity observed during these periods might not solely represent information of the rat position. So, the rat occupation over time in the maze is non-uniform and this needs to be reflected back to the LSTM model training. Wihtout this consideration, the LSTM or any other statistical decoder will favor the decoding result toward those points that the rat spends most of its times. This looks an unbalanced learning problem for position decoding, and it should be compensated in the training step. There are multiple solutions to address this, first including a non-uniform sampling over time. Second, defining the cost function as a function of velocity or other information which compensates the non-uniform occupancy. We can also address the unbalanced occupancy problem by adding a weight to the learning mechanism. We define the training objective function by a weighted Mean Squared Error (MSE) where the weighted is a function of the velocity. The weighted MSE is define by

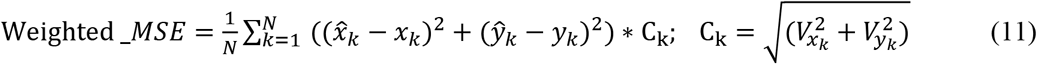

where *N* is the total number of samples in a training batch, (*x_k_*, *y_k_*) is the rat position at the time interval k, and 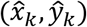 is the estimated rat position. (*V_xk_*, *V_yk_*) is the velocity of the rat at the time-step *k*, where the weight is defined by the amplitude of the velocity. We use this cost function to train the position-estimator layer. This weighted cost function debilitates effect of unbalanced data from the regions that rat spends more time in and helps to smooth effects of biased dataset. Regarding to other layers in the LSTM topologies, the velocity-estimator layer train by MSE cost function because it solves a regression problem. The geometry-estimator layer which extracts probability of the rat appearance in a area of the maze, train by a cross-entropy cost function which is a well-known cost function for a multinomial classification [45].

In the next section, we use the training algorithm to update each of propose models paramaters (numerical exact solution, approximate filter solution, and three LSTM network topologies) for decoding a rat’s movement trajectory in 2-D spaces.

## 3 Application of SSPP and LSTM approaches in spiking activity of hippocampus place cells ensemble

In this section, we start with describing the experiment and the neural data recorded from a rat hippocampus while moving in a W-maze. We then discuss different metrics to assess accuracy of different solutions. Finally, we configure the LSTM and SSPP models introduced in previous sections to estimate the rat movement trajectory from the cell ensemble spiking activity properly.

### 3.1 Data Description

The neural data were recorded from 62 place cells in the CA1 and CA2 regions of the hippocampus brain area of a Long-Evans rat, aged approximately 6 months. The rat has been trained to traverse between the home box and the outer arms to receive a liquid reward (condensed milk) at the reward locations. **Figure 4** shows the maze structure and the rat's movement trajectory in 2-D spaces, where the rat position at each time step is represented by (*x*, *y*) coordinates. Spiking activity of these 62 units were detected offline by choosing events that their peak-to-peak amplitudes were above a threshold of 80uV in at least one of the tetrode channels [46]. In the experiment process, the actual rat’s position was measured by a video tracking software which was used as the ground truth for the position. We used a 15 minutes experiment session, sampled with a time resolution of 33 milliseconds, to train and then evaluate the models performance. We divide the data to training and test datasets. We used 85% of the data, which corresponds to 23209 data points to train the LSTM networks and SSPP models. The remaining15% of the data, corresponding to 4095 data points, were used to test the trained models estimation result.

**Figure 4:**
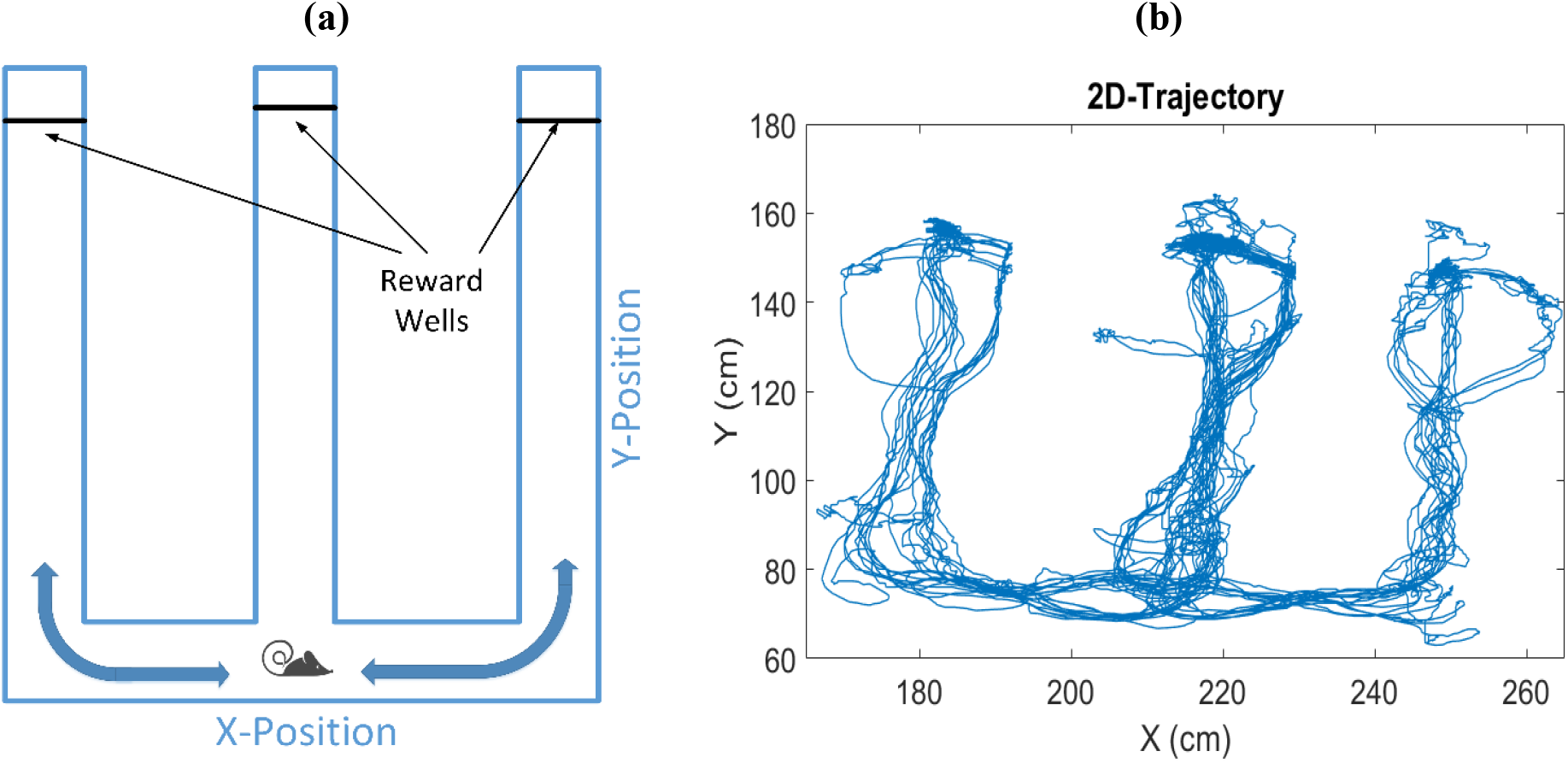
The maze structure and the rat's movement trajectory. **a.** Structure of the experiment, W-maze. The rat moves from the center arm to the left and right arms to get food rewards. **b.** Movement trajectory in X and Y directions during the experiment

## 3.2 Performance Metrics

To analyze the accuracy of the approaches, we calculated both the mean of L1-norm, called least absolute errors (LAE), and the root mean square error (rMSE) [47] between the rat actual position and the estimated position. LAE and rMSE performance metrics are defined by:

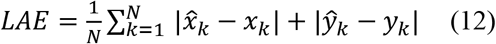

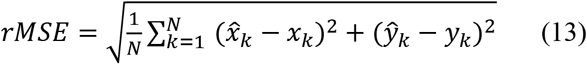

where *N* is the total number of timesteps, (*x_k_*, *y_k_*) is the rat actual position at timestep k, and 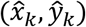 is the decoder estimation of the rat position. There are unique properties of these metrics which make them good choices as performance metrics for this problem; lower rMSE and LAE show a more accurate decoding result. But, rMSE is more sensitive to larger error (larger than one). The rMSE measure effects the error by powering it by two. In other hand, in LAE measure, error contributes equally in the overall decoding performance. For the SSPP models, we can also build other metrics based on the decoder posterior distribution, like 95% HPD coverage area which is the 95% highest probability density region of the computed posterior distribution of the rat’s position [42].

### 3.3 Parameters setting in DNN and SSPP

In this section, we setup our proposed LSTM and SSPP models to fit to the data. For the SSPP models, we assume the rat movement follows a random walk in 2-D spaces, we set *A_k_* = *diag*[1,1] and *B_k_* = [0,0]^*T*^ defined in equation(5), where *Q* (equation (5)) is a diagonal covariance matrix defined by

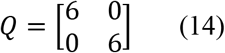

Where its terms extracted from the rat movemnts statistics during the training session. For the Numerical exact solution, The spatial resolution in both the *x* and *y* dimensions is 2 cm; given the maze dimension, we have 50 samples in the *x* direction and 58 samples in the *y* direction. We draw 4000 samples at each spike time to calculate the likelihood function and rat position posterior estimation using Riemann sum integral method [48]. For the approximate filter solution, discussed in Appendix A, we set *α_m_* = 0.001 and *α_d_* = 0.001 to calculate likelihood function and the rat posterior.

In this experiment, the rat’s position is constrained to the maze area. So, the rat’s movement is bounded by the maze, and the formuation for SSPP models (section 2.1), which defined for the rat movement inside the maze, may be misspecified. To address this issue, we add a penalty term to the likelihood function of SSPP models that accounts for the toplogy of the maze [23]. The revised prediction process is expressed as

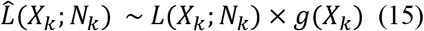

where *g*(*x_k_*) is close to zero for x-y coordinates outside the maze area and one otherwise.

In LSTM topologies, we use weighted MSE on (equation (18)) as the position-estimator layer which consists of four LSTM nodes (cost function *L_1_*). We use Traditional MSE cost function for the velocity-estimator layer which is consist of two LSTM nodes (cost function *L_2_*). As we discussed (section 2.2.1), we should segment the experiment area which the rat moves. **Figure 5** shows suggested segmentation approach for encoding the experiment area for training the geometry-estimator layer. A further explanantion of the specific segementaityon of the regions can be found in the Appendix B. This layer consists of four LSTM nodes for extracting probability of all four segements and uses cross-entropy cost function (cost function *L_3_*)[45].

**Figure 5:**
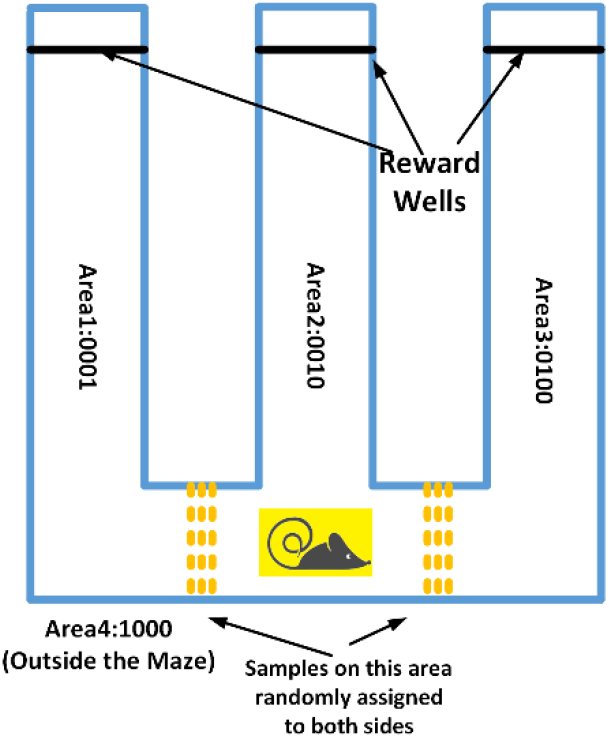
Segmentation mechanism for the maze structure. Three arms plus outside area encoded as Area1 to Area4. The areas between two adjacent arms assigned randomly to one of these near areas.

To perform the learning process, we try to minimize total cost function defined by *L* = *L_1_* + *L_2_* + *L_3_* using RmsProp optimizer [49], with a learning rate of **0.001**, a decay factor of 1*e*^−6^ for the first 100 iterations, and a learning rate of **0.0001** for the 100 next iterations, which is the same for all topologies in training section.

## 4 Results

We analyzed the decoding performance of the LSTM and SSPP models in decoding 2-D movement trajectory of the rat using place cells’ spiking activitiy. We examined the metrics described in section 3.2, and the computational cost (i.e. computational time) of the decoer models. **Table 2** shows the performance of the SSPP and LSTM models. The performance result of the approximate filter solution shows that even though the accuracy drops only about 15%, 25%, and 2%, in LAE, rMSE, and 95% HPD measures, respectively, as the **Table 2** shows the computation time in approximate filter solution is much less than the Numerical exact solution (the approximate filter solution is about 2780 times faster than the Numerical exact solution). Beside that, the performance of different LSTM network topologies show that our a assumption in utilizing higher order elements of movement including movement speed, and direction of movemet in the hierarchical LSTM networks, improves accuracy of the network in decoding 2-D position from spiking activity.

**Table 2:**
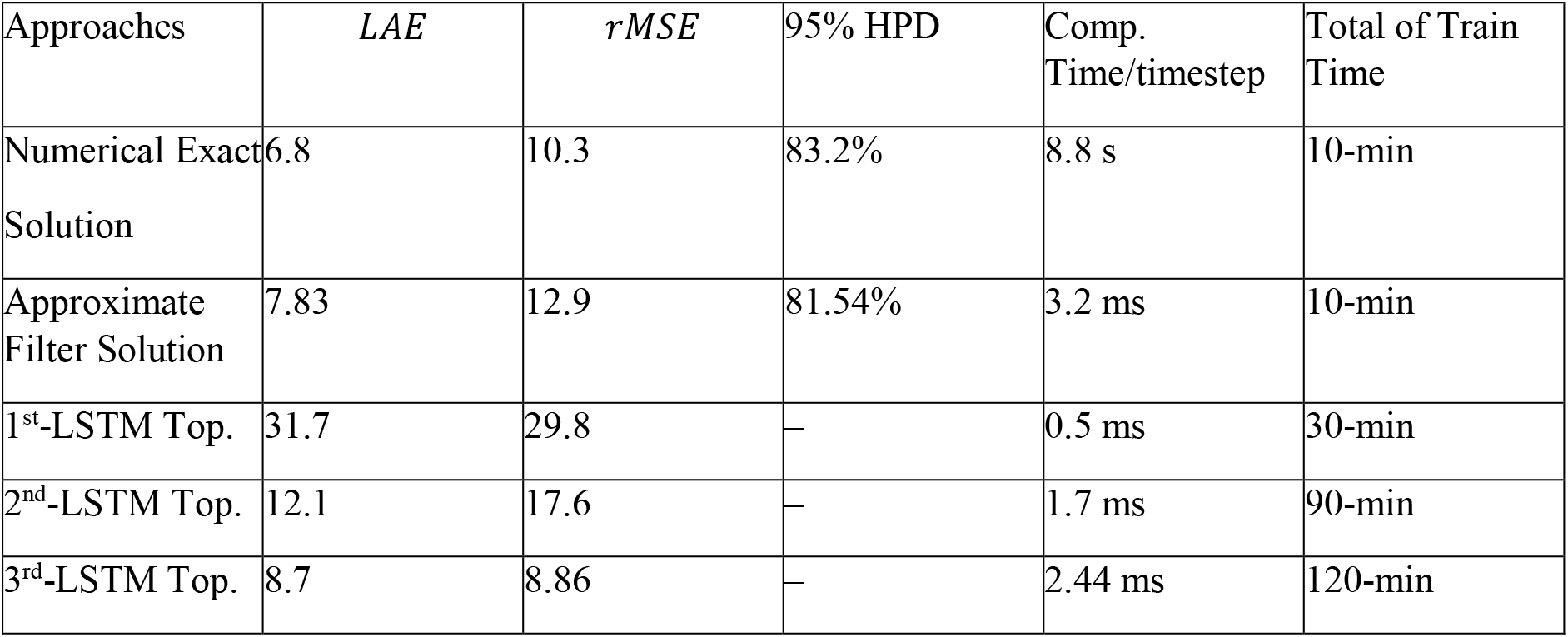
Performance and computational cost result.

| Approaches | *LAE* | *rMSE* | 95% HPD | Comp. Time/timestep | Total of Train Time |
| --- | --- | --- | --- | --- | --- |
| Numerical Exact Solution | 6.8 | 10.3 | 83.2% | 8.8 s | 10-min |
| Approximate Filter Solution | 7.83 | 12.9 | 81.54% | 3.2 ms | 10-min |
| 1<sup>st</sup>-LSTM Top. | 31.7 | 29.8 | — | 0.5 ms | 30-min |
| 2<sup>nd</sup>-LSTM Top. | 12.1 | 17.6 | — | 1.7 ms | 90-min |
| 3<sup>rd</sup>-LSTM Top. | 8.7 | 8.86 | — | 2.44 ms | 120-min |

The 3^rd^ LSTM network topology has the best accuracy among all three LSTM network topologies in decoding the rat position. Although its accuracy drops less than 13% in LAE metric compared to the Numerical exact model, but the 3^rd^ LSTM network topology is 3600 times faster than the exact solution.

In summary, the 3^rd^ LSTM network topology and approximate filter solution are the best among the models based on computational time and accuracy. **Figure 6** shows the mean of estimated movement trajectory for the approximate Filter solution, the 3^rd^ LSTM network topology, and the Numerical exact solution.

**Figure 6:**
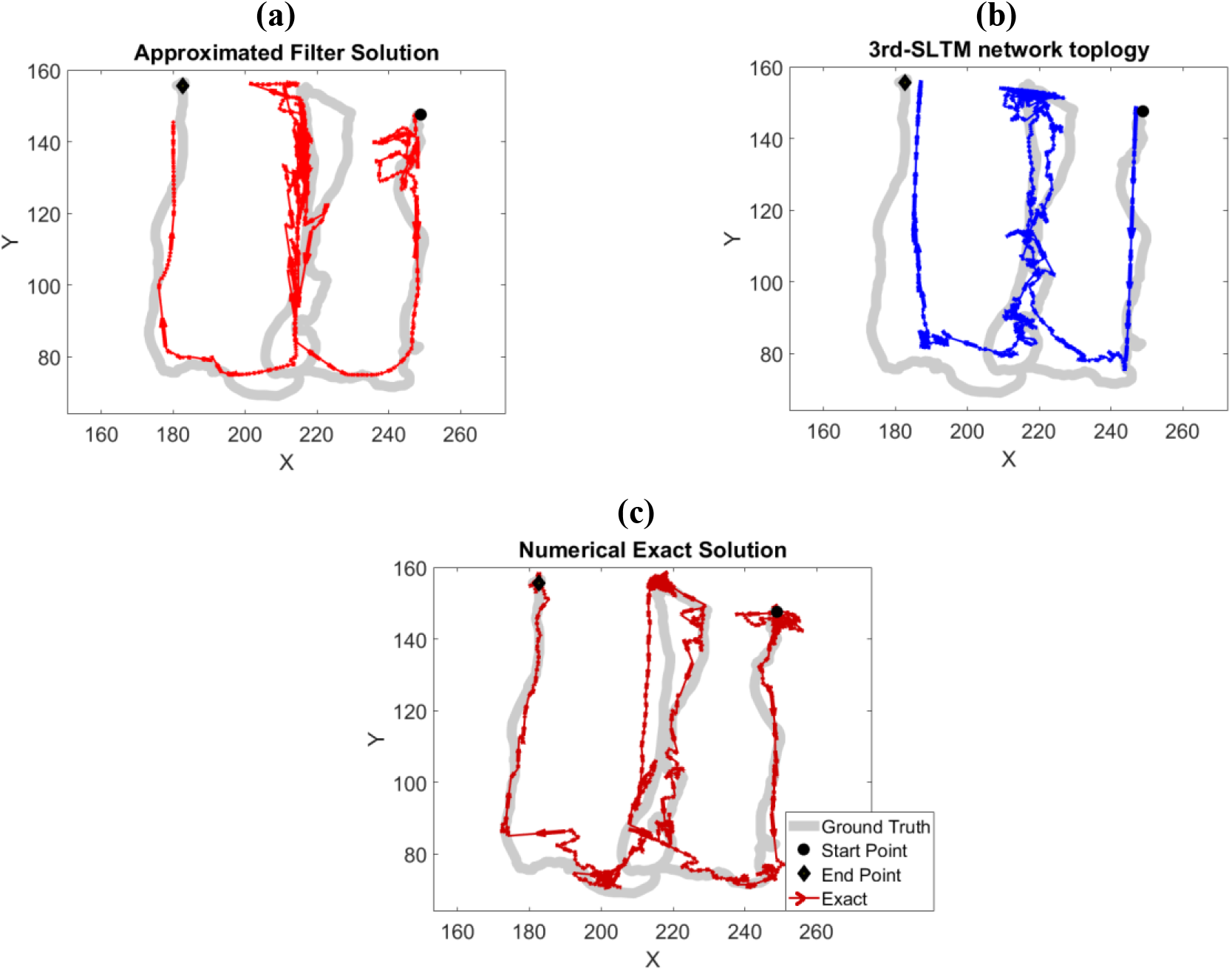
**a**. Mean of estimated trajectory in approximate filter solution. **b**. Mean of estimated trajectory in the 3^rd^ LSTM network topology. **c**. Mean of estimated trajectory in Numerical exact solution.

We already know SSPP models encode the place cell’s area of activity (receptive field) of indivudal neurons [50, 51], which characterize the realatinship btween a cell firing activity and positionis. In the next section, we study how information about receptive field is being captured in different LSTM models.

### 4.1 Analysis Result of the Encoding Mechanisms

Each place cell has a distinct receptive field; the receptive filed of a place cell corresponds to areas in the space where the cell firing activity grows significantly from its normal firing rate. Place cells' receptive fields might have different shapes and also cover different areas of the maze; generally, the receptive field of a place cell might be multi-modal and change over time as well. **Figure 7** shows the spiking activity of different cells as the rat traverse the maze. Receptive field of these samples cells are shown in **Figure 7**. In this analysis, we choose an input channel of the decoder model which is assigned to a specific place cell. Then we replace its spiking activity with a periodic spiking activity with a firing rate close to its natural peak firing rate. The period of the spiking patterns has been chosen carefully based on the spiking rate of the place cell; note that other place cells have their recorded spiking activity. With this set up, we then decode the rat movement trajectory based on new data.

**Figure 7:**
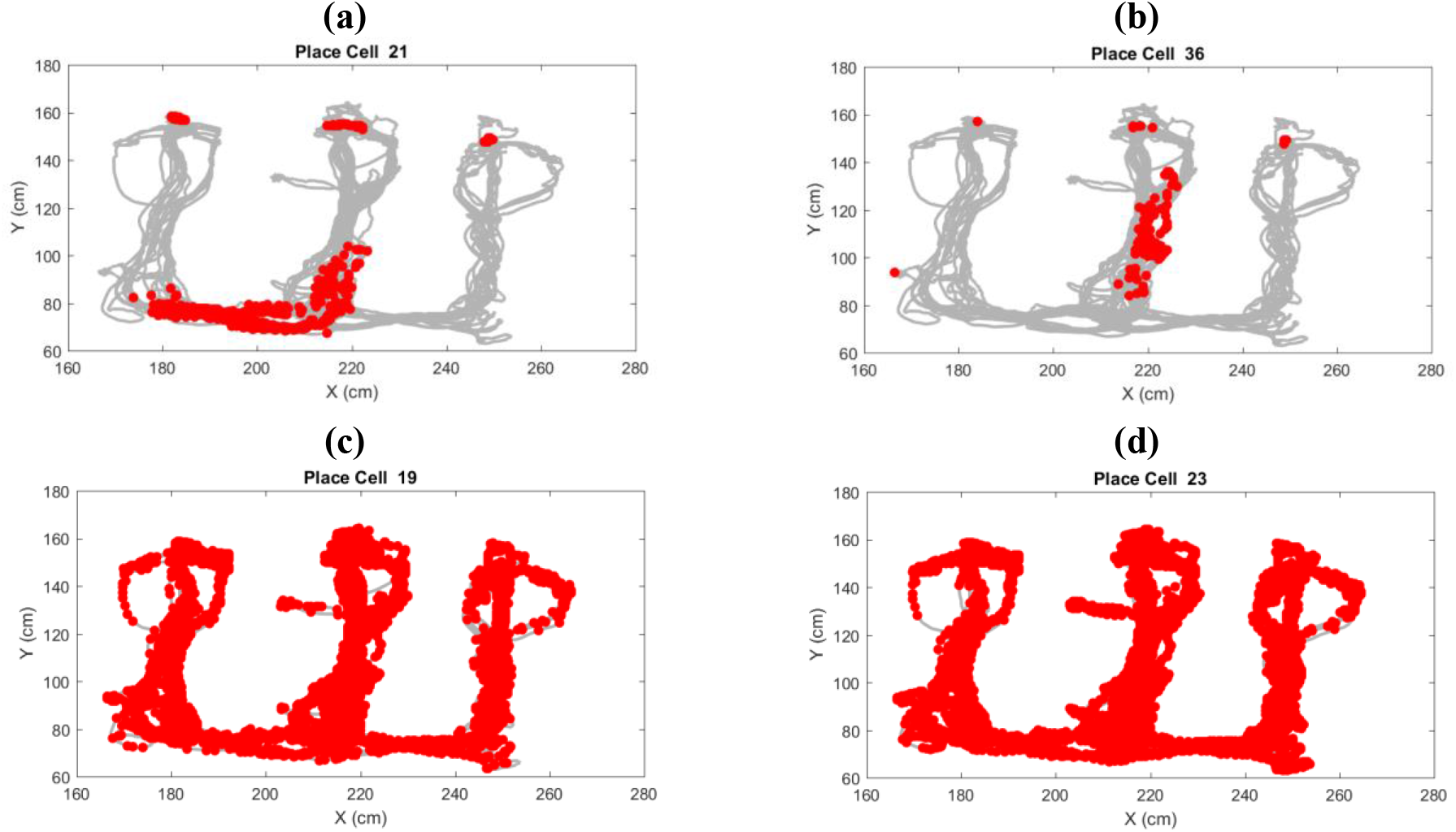
The spiking patterns of two different groups of place cells with a localized receptive field and putative place cells with a less distinct receptive field. **a-b**. place cells with a distinct receptive field. **c-d**. place cells without a localized receptive field.

The hypothesis is that if the decoder model is capable of capturing information of receptive field for each place cell, the decoder should decode a trajectory that converges to the corresponding cell receptive field area; no matter where the decoder starts in the maze. In the other hand, if the corresponding place cell does not have a distinct receptive field, the decoded trajectories should stand near starting point or move to the different points over the maze.

For place cells with a distinct receptive field. We replace a cell spiking activity with its natural peak firing rate pattern individually; then we run the decoder from three different initial points over the maze area. As the result show (fourth and fifth rows of **Figure 8)**the movement trajectories given by the 3^rd^ topology of the LSTM network and the approximate filter solution converged to these receptive field areas. The result indicate these models captured the information of place cells’ receptive field with a distinct area. However, the movement trajectories given by the 1^st^ and 2^nd^ topology of the LSTM networks did not converge to the receptive field areas (second and third rows of **Figure 8).**

**Figure 8:**
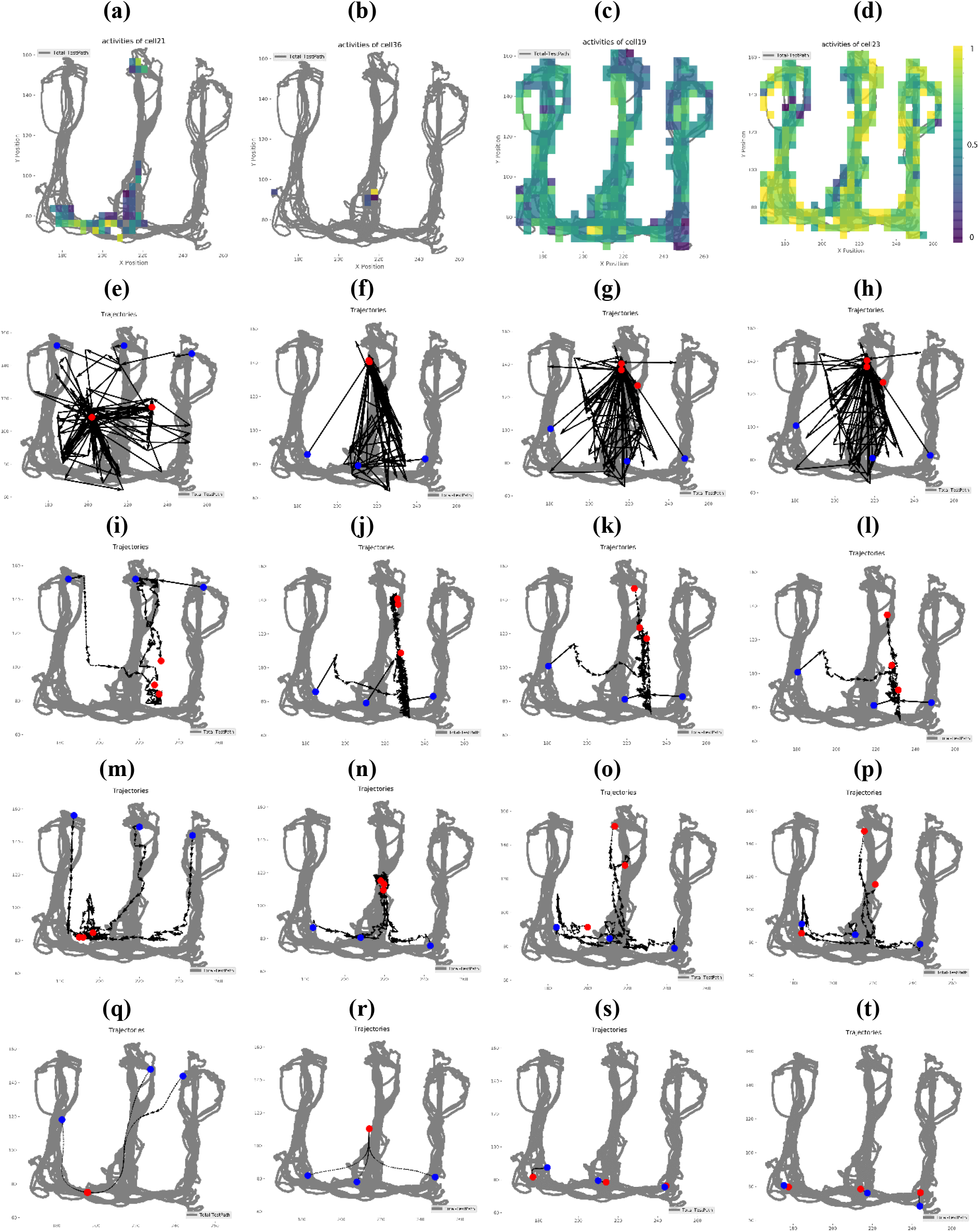
**a-d**. Spiking activity density over the maze surface for place cells 21, 36, 19, and 21, respectively. **e-h**. Decoding result of the 1^st^ LSTM network topology by considering a periodic stimulus replaced the place cells spiking activity. **i-l**. Decoding result of the 2^nd^ LSTM network topology by considering a periodic stimulus replaced the place cells spiking activity. **m-p**. Decoding result of the 3^rd^ LSTM network topology by considering a periodic stimulus replaced the place cells spiking activity. **q-t**. Decoding result of the approximate filter solution by considering a periodic stimulus replaced the place cells spiking activity.

For place cells without a distinct receptive field; the result show (fourth and fifth rows of the **Figure 8)**the 3^rd^ LSTM network topology and the approximate filter solution decoded movement trajectory neither moves nor converges to a specific area in the maze, it means the 3^rd^ LSTM model and the approximate filter solution captured the information of place cells’ receptive fields without any distinctive area too. In other hand, the 1^st^ and 2^nd^ LSTM network topologies did not decode movement trajectories which contradict the hypothesis (second and third rows of **Figure 8)**. In other word, the 1^st^ and 2^nd^ LSTM network topologies neither captured the information of place cells with a distinctive receptive field area nor the information of place cells without any distinctive receptive field area.

The results show the approximate filter solution and the 3^rd^ LSTM network topology have similar behavior with spiking activity data comapare to the exact solution which is a gold strandard for this problem even though they are built using completely different modeling structures.

## 5 Discussion

In this research we porposed LSTM and SSPP models to decode a rat movement in 2-D spaces from its spiking activites. We studied two different metrics - *LAE*, *MSE* and computational efficiency - to compare the performance of DNN and SSPP models in 2-D spaces decoding problem. The result show that the approximate Filter solution and 3^rd^ LSTM network topology significantly reduce computational time of the decoding step; compared to the Numerical exact solution. These models are more than 2000 times faster than the Numerical exact solution with negligible decrease in prediction accuracy accuracy. To design these models we faced multiple challenges which our methods to address the discussed in bellow.

LSTM networks are flexible and powerful tools to extract information from complex and multi-dimensional time-series data including the neural activity. The idea of using LSTM network topologies to characterize neural activity has a great promise in the neural data analysis and in particular neural decoding problems. However designing a proper LSTM network topology is challenging. The first challenge was interperatation ability of the LSTM network. To address that, in the 3^rd^ LSTM network toplogy we extracted higher order information of the rat movement (like velocity, movement direction, and geometry of the maze structure) in a specific hierical structure and use them to decode the rat position in the maze. By doing this we increased the interperatation ability and accuracy of LSTM network topologies. This leaded to a LSTM network topology with better performance and generalizability in the neural decoding task. Our approach in developing this topology may helps other researchers to build their specific LSTM network topologies for other neuroscience data.

The other challenge was identifying the size of the LSTM network topology, which indicates the number of free parameters. As the size of the LSTM network increases, the training data must also be increased to prevent overfitting problem. In this research by using right information (geometrical features and velocity) and embeding them in right hierarical structure, we could maintained the number of the LSTM topologies parameters in order of 10^3^. These topologies are much smaller than other networks previously used for neural decoding problems [19], which are designed blindly and without considering the characteristics of the problem that they designed the LSTM network for it. So, there is no need for a large training dataset to properly train these models.

The other challenge was overfitting issue, to address that we used to reduce the effect of overfitting in traning the LSTM topologies is technique [52]. In each layer of our LSTM network topologies we set the drop out ratio to 0.2, meaning that 20 percent of parameters in each layer are randomly chosen to be ignored in each iteration of training. This technique improves the LSTM network topologies generalizability and reduce effect of overfitting on the training data [53].

We addressed a couple of challenges in designing a proper LSTM network topology fro neural decoding problems, but there are still doubts about how the trained networks perform with unseen place cells’ receptive filed information. The 3^rd^ LSTM network topology can extract features of each place cells’ receptive fields, but we don’t know if it combines them, or extracts collective behavioral features of place cells’ receptive fields to estimate the rat position. A promising solution to address this is computing a posterior for the network parameters. We can use the Bayesian approach combined with the learning process to calculate this posterior for the network parameters, which is our concern for future work [54].

Our analysis shows that the approximate filter solution is a proper approximation for the Numerical excat solution in this particular neuroscience experiment, but to design it, we also faced a couple of challenges. The first challenge was computational efficiency, by selecting small values for *α_d_* and *α_m_* parameters, even though we can get better accuracy in decoding the movement trajectory, this led to increase in the size of MoG which process in each time-step and this means increase in computational time. So, it is a tradeoff between gainig accuracy and computational effieciency. We used the parameter set based on its effect on accuracy and computational effiecieny studied in [35]. A better solution for selecting *α_d_* and *α_m_* parameters is a adapting solution. Means updating values for these parameters in each time step based on our observation of spiking activity and the postieror frequently, which our concern for our future work.

The second challenge was, the primary hypothesis in this approach is that we modeled CIF of each neuron with a MoG, which means by considering large enough MoG we can model any CIF of neourons, but due to the size of MoG, it might debilitate computationa the efficiency of the approximate filter solution. So, it is crucial to select the right MoG to modle each neuron’s CIF. Here we choosed a MoG (set of 30 gaussian models) to cover the hole are of the maze which enables the model to cover all areas of the maze for estimating each neuron’s CIF and maintain it size small enough to avoid computationl complexity. Finally, as we discussed in the application section to compensate misspecification of the rat movement outside the maze structure, we added a penalty term to the likelihood function of SSPP models that accounts for the maze toplogy. This penalty term push the likelihood towards the area inside the maze and avoid any misspecified position autside the maze areas.

## 6 Conclusions

In a context where a comprehensive study which systematically assess attributes of DNN and SSPP approaches on a common neuroscience data analysis problem is lacking we have presented a rational approach to design neural decoder models. We compared the methods in common neuroscience problem. for the DNN models, we presented an effective way in embedding higher order information like velocity and maze structure in the LSTM networks for gaining accuracy and generalizability for a class of point-process filter problems. In the SSPP approach, we developed an approximate filter solution for this class of filter problems. We used these solutions in decoding 2-D position from spiking activities of a population of neurons in the hippocampus area of a rat, when it’s navigating through a W-shaped maze. The approximate filter solution and LSTM networks are more than 2000 times faster than the Numerical exact method with insignificant decrease in position decoding accuracy. We also investigated both modeling approaches mechanisms in encoding place cells' receptive field information. As the result shows, the approximate filter solution and the 3^rd^ LSTM network topology can properly capture the activities of place cell and encode each receptive field, even though they have completely different natures and modeling structures.

## 7 Acknowledgments

This research was partially funded by R01 MH105174 and SCGB grant #320135. A copy of the source code utilized in this research along with sample data can be found in the GitHub link^1^.

## APPENDIX A: Dropping and Merging Procedure of a GMM

In the dropping process, we take one of the *P* components out and rescale other mixtures’ weight to keep their sum equal to one. We then check the distance between these mixture models *Q* and *P* to find which mixture model might be dropped [35]. We repeat this procedure until a stopping criterion is met. The dropping process is as follows:

**Table 1.**
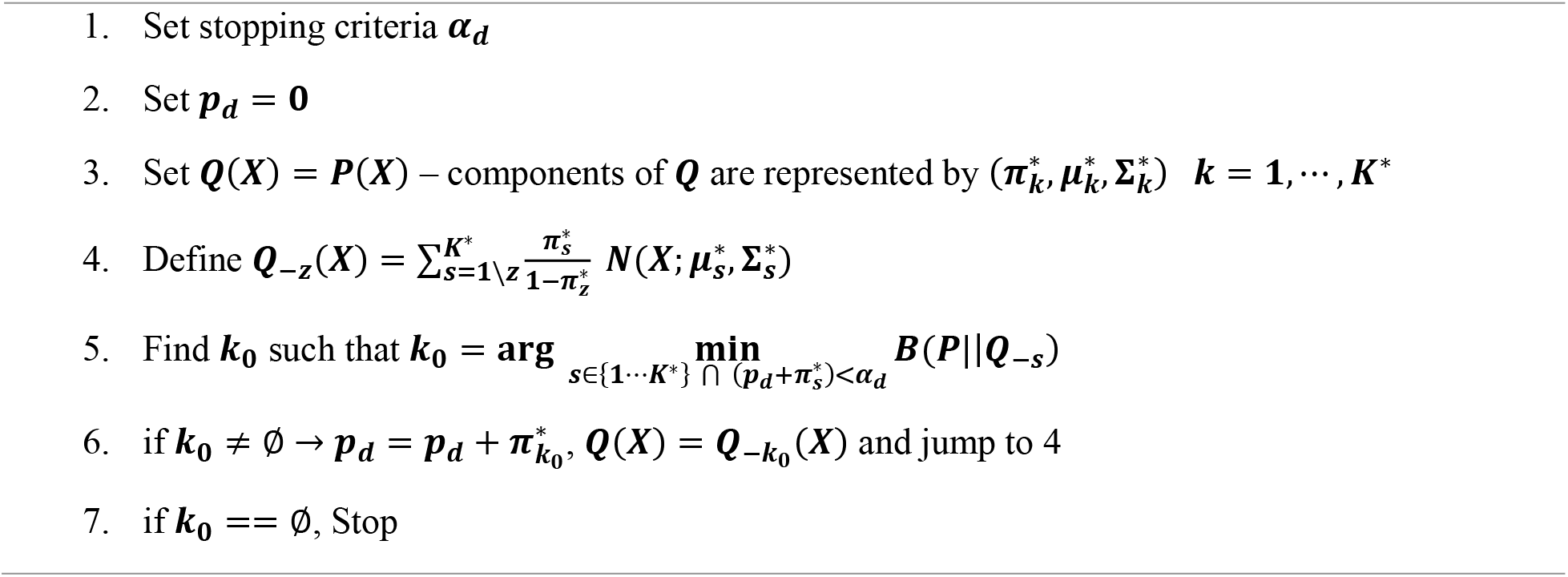
Dropping Process Algorithm

In the merging process, we search for a pair of mixture components which can merged while the growth of *B*(*P* ∥ *Q_i∘j_*) is minimal – *Q_i∘j_* represent the new mixture model with its *i* and *j* mixture components being merged. The merging process is run sequentially; thus, per each iteration, number of *Q* drops by one. The merging process is repeated until a stopping criterion is met. The merging process is as follows:

**Table 2.**
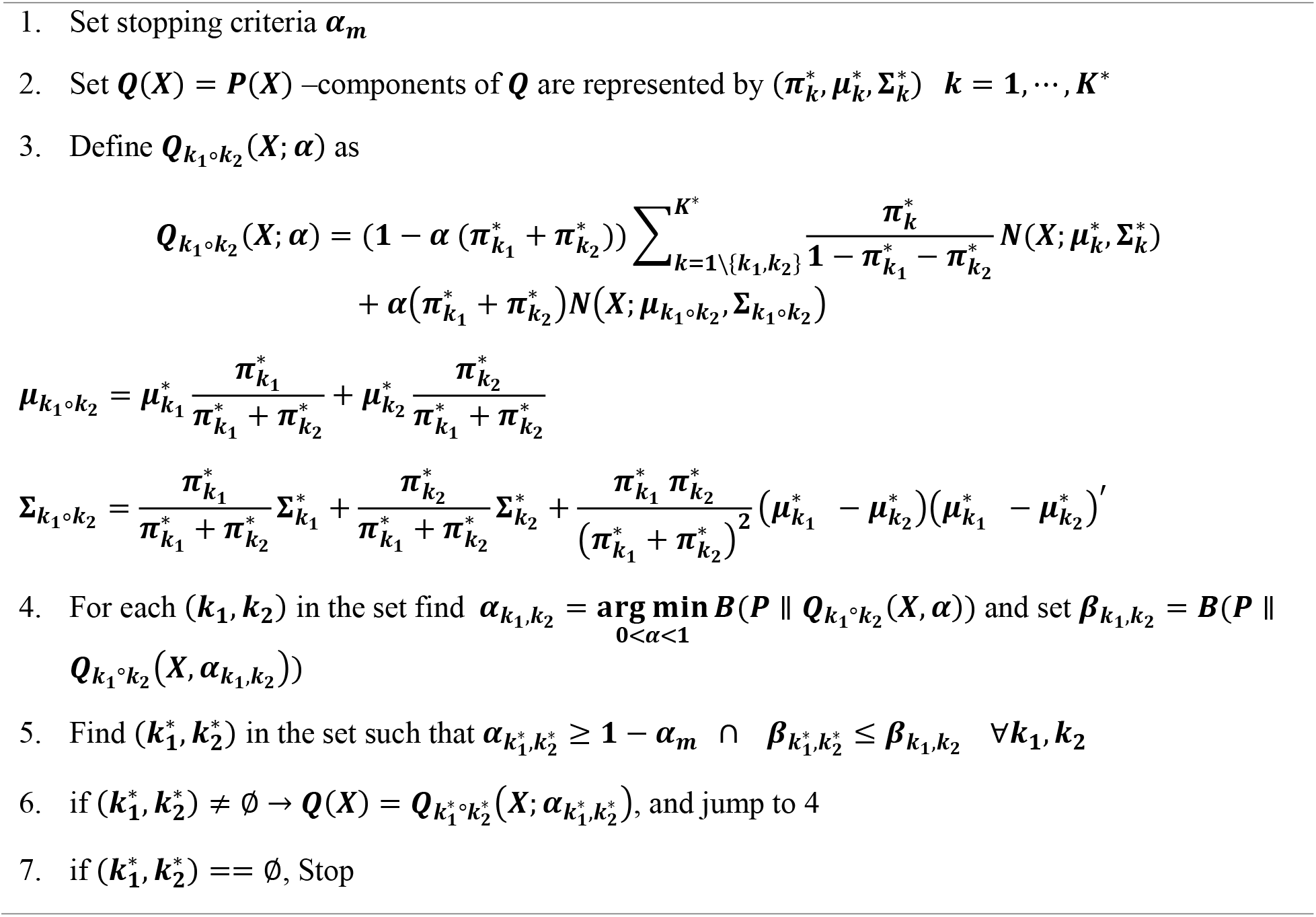
Merging Process Algorithm

## Footnotes

1 https://github.com/mrezarezaei/Decoding_position_and_velocity_from_spiking_activity

## Notes

### Competing Interest Statement

The authors have declared no competing interest.

